# Short-term dietary change rapidly remodels microbial community assemblages and reprogrammes systemic immune phenotypes

**DOI:** 10.64898/2026.07.16.738828

**Authors:** Rebecca C. Simpson, Matthew McKay, Miguel Castañeda, Mirei Okada, Reem Elnour, Zhen Bao, Jian Tan, Mark Molloy, Georgina V. Long, Erin R. Shanahan

## Abstract

Diet is a major determinant of the gut microbiome and immune homeostasis, yet the extent to which short-term dietary interventions can remodel established microbial communities and reprogramme immune phenotypes following long-term western diet consumption remains poorly understood. Here, we investigated temporal dynamics of the gut microbiome, microbial metabolites, intestinal barrier function and local and systemic immune responses following diet switching. Mice were fed either a standard chow or a western diet for 8 weeks before remaining on these diets or switching to the alternate diet for 2 or 4 weeks. Long term consumption of chow and western diets resulted in distinct gut microbial communities and differences in intestinal permeability. Diet switching rapidly remodelled microbial community structure within two weeks, with substantial bidirectional changes in community composition. Despite these changes, the relative abundance of several taxa remained influenced by prior dietary exposure. In contrast, faecal SCFA profiles remained largely associated with long-term diet, indicating that microbial metabolic outputs were altered more slowly than microbial community composition. Mass cytometry revealed progressive remodelling of local (MLN) and systemic (PBMC and spleen) immune responses following dietary switching. Activation-associated immune phenotypes, including Ki67+ and PD-1+ B and T cells, inflammatory monocytes and RORγt+ regulatory T cells, rapidly responded to diet switching, whereas overall B cells, regulatory T cells and effector memory T cells retained signatures of long-term dietary exposure. Together, these findings demonstrate distinct temporal dynamics across the diet-microbiome-immune axis, whereby gut microbial composition and immune activation states remain highly plastic, while microbial metabolic outputs and several memory and regulatory immune phenotypes exhibit persistent dietary imprinting. These findings highlight the potential utility of short-term dietary interventions to modulate host-microbiome interactions and immune homeostasis.

## Introduction

Diet and the gut microbiota are important factors that regulate immune homeostasis both at the gut epithelium and systemically. Long-term dietary patterns strongly shape the composition and function of the gut microbiota and influence the capacity of the microbiota to maintain intestinal barrier integrity and modulate immune function^1–4^. A prominent way through which microbes modulate systemic immunity is via the production of metabolites. This is dependent on both the type of microbes present in the gut as well as the nutrients ingested and utilised by these microbes^5,6^. These metabolites can alter immune activity by direct signalling, such as through metabolite sensing G-protein coupled receptors (GPCRs) or indirectly through regulating gut barrier integrity and intestinal permeability, thus altering immune cell exposure to microbes and microbial products^7,8^. Suboptimal diets such as the consumption of a Western diet characterised by low levels of fibre consumption and high intake of saturated fats and sugar have been linked with a loss of beneficial gut microbes, a reduction in SCFA levels and reduced gut barrier integrity, leading to increased inflammation both locally in the gut and systemically^9–14^. Unsurprisingly, it has therefore been implicated in the development of a variety of metabolic and inflammatory diseases^4,8,15^. In contrast, a fibre-rich diet has been shown to be protective against the development of colitis, colorectal cancer as well as allergies^16–19^. There is increasing interest in understanding how rapidly dietary interventions can remodel the microbiota-immune axis in clinical settings where immune function must be modulated over relatively short timeframes, such as during cancer treatment. Dietary modification therefore represents a promising strategy to therapeutically target the microbiota-immune axis across a variety of disease contexts.

Although the detrimental effects of a long-term western diet on the gut microbiota and immune system are well established, considerably less is known about microbiome-immune dynamics following a short-term dietary switch and the capacity to reprogramme long-term immune priming. While the gut microbiota responds rapidly to alterations in diet, it remains unclear whether short-term dietary intervention can similarly remodel immune phenotypes established during prolonged dietary conditioning. This distinction is particularly important because immune phenotypes may exhibit different response kinetics to dietary intervention than the gut microbiota, potentially limiting the efficacy of short-term dietary interventions. We therefore used a standard chow versus a western diet to model the extremes of a healthy balanced versus an unbalanced diet and assessed the capacity of a short-term dietary switch following the long-term priming diet to alter established microbial community assemblages and reprogramme local and systemic immune responses. Here, we show that short-term dietary switching rapidly remodels gut microbial community structure and induces substantial changes in local and systemic immune phenotypes. However, while many immune populations rapidly adapt to the new dietary environment, others retain signatures of long-term dietary conditioning, demonstrating distinct temporal dynamics across the diet-microbiome-immune axis.

## Results

### Western diet alters gut microbial community composition

Mice were fed either a standard chow (CD) or a western diet (WD) for 8 weeks before remaining on these diets or switching to the alternate diet for 2 or 4 weeks (Figure 1A). This resulted in four experimental diet groups: long-term chow (CD), long-term chow to short-term WD (CD>WD), long-term WD (WD) and long-term WD to short-term chow (WD>CD) (diet composition summarised in Supplementary Figure 1A). As expected, the WD-fed mice gained weight more rapidly than the CD-fed mice, which was reversed with the switch to CD (Supplementary Figure 1B-D). To characterise longitudinal changes in the gut microbiota, faecal samples were collected throughout the study and analysed using 16S rRNA gene amplicon sequencing. Following the initial 8 weeks on the respective long-term diets pripr to the diet switch (Day 61), alpha diversity did not differ between CD and WD-fed mice assessed by the Inverse Simpson’s or Shannon indexes, however species richness was significantly greater in mice on chow (p = 0.0280) (Figure 1B, Supplementary Figure 1E-G). In contrast, beta diversity analysis using Bray-Curtis demonstrated distinct microbial community composition between CD and WD-fed mice (PERMANOVA, p= 0.001) (Figure 1C, Supplementary Figure 1K). Greater within-group variability was observed among CD-fed mice, although this accounted for only ∼11% of the variance compared with ∼54% for the between-group comparison of CD and WD microbiomes.

**Figure 1.**
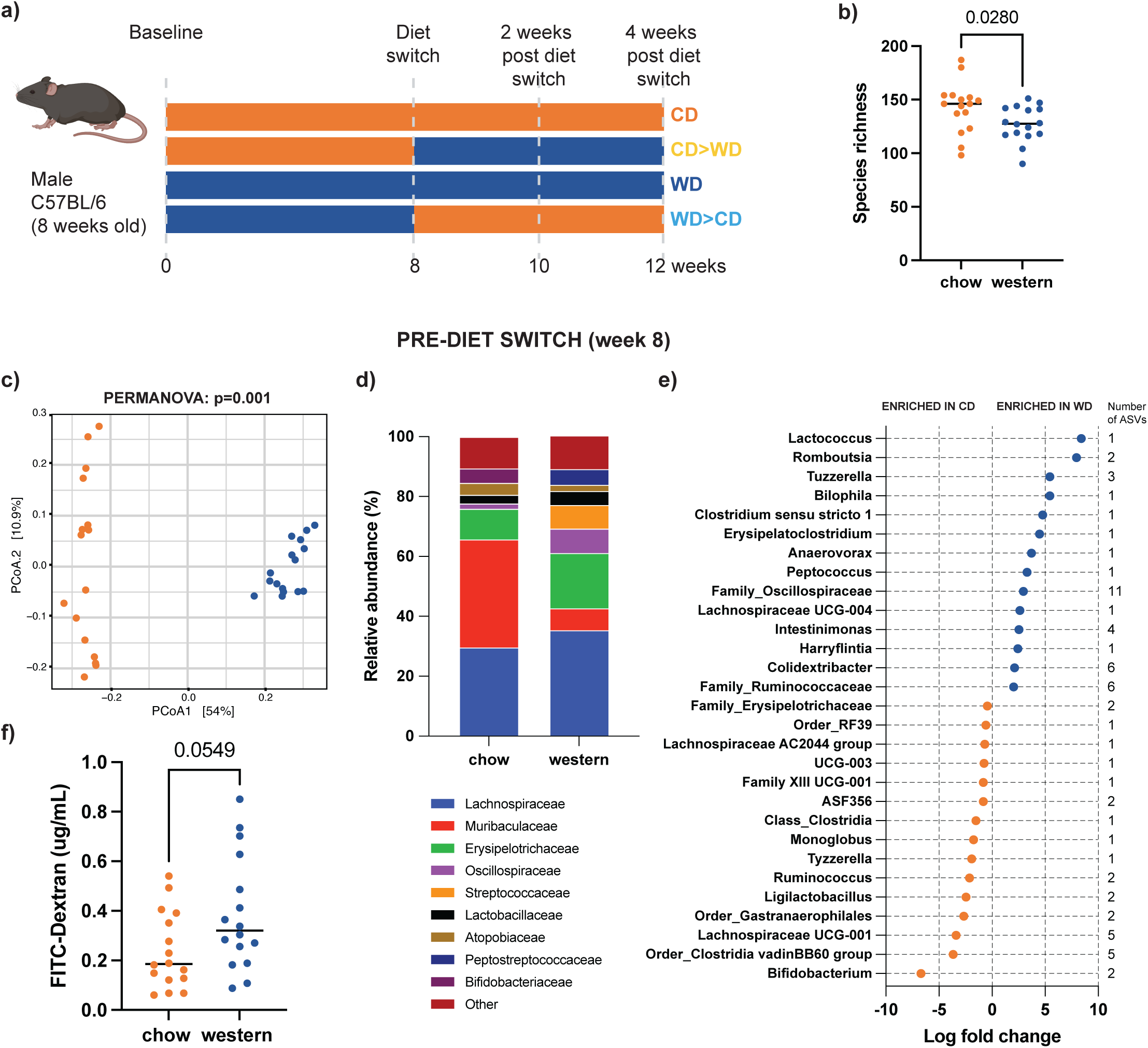
Distinct gut microbiome profiles are associated with long-term western and chow diets. (a) Schematic overview of mouse study. Male C57BL/6 mice were fed either a chow or western diet for 8 weeks (n=32) before remaining on these diets or switching to the alternate diet for 2 or 4 weeks, resulting in 4 diets groups: long term chow (n=8), long term chow to short term western (chow>western)(n=8), long term western (n=8) and long-term western to short term chow (n=8). Faecal samples were collected from all mice after 8 weeks on diets prior to the diet switch and profiled by 16S rRNA gene amplicon sequencing. (b) Alpha diversity calculated by species richness grouped by long-term diet, chow (orange) (n=16), western (blue)(n=16). (c) Principal coordinate analysis (PCoA) of beta diversity of microbiome samples calculated by Bray-Curtis dissimilarity, coloured according to long-term diet. P-values were calculated using PERMANOVA. (d) Family level composition of mice on the chow verse western diet, average relative abundance (10 most abundant families overall) plotted. (e) Differential abundance at genus level assessed using ANCOM-BC. Significant differentially abundant taxa (q < 0.05) are presented as log2 fold change between mice on the chow and western diets. (f) Pre-diet switch intestinal permeability split by long-term diet. Dots indicate individual mice, bars indicate median. P-values calculated by Mann-Whitney U rank test.

Differences in the composition of the gut microbiota based on long-term diet are evident when comparing the most abundant family level taxa across all mice (Figure 1D). Differential abundance analysis using ANCOM-BC at the genus level, identified 29 genera that significantly differed between CD and WD mice (Figure 1E). Mice on the western diet were enriched in *Lactococcus, Romboutsia, Tuzzerella, Bilophila, Clostridium sensu stricto 1, Erysipelatoclostridium* and *Anaerovorax,* whereas mice on the chow diet were enriched in *Bifidobacterium, Clostridia vadinBB60 group*, *Lachnospiraceae UCG-001, Ligilactobacillus* and *Ruminococcus*. To establish whether long-term dietary exposure was also associated with altered intestinal barrier function prior to dietary switching, intestinal permeability was assessed using the *in vivo* FITC-dextran assay. After 8 weeks of dietary intervention, WD-fed mice exhibited a trend towards increased intestinal permeability compared with CD-fed mice (p = 0.0549; Figure 1F).

### Short-term diet switch rapidly alters gut microbial community composition

We next assessed how the gut microbiota of mice changed in response to the diet switch by comparing faecal samples collected immediately prior to dietary switching and endpoint caecal contents (Figure 2A). Alpha diversity remained largely unchanged following dietary switching; however, mice switched from the western diet to chow (WD>CD) exhibited a significant increase in species richness, whereas a trend towards a reduction in species richness was observed with the switch from chow to the western diet (CD>WD) (Supplementary Figure H-J). Bray-Curtis dissimilarity demonstrated that microbial community composition clustered primarily according to the diet at the time of sampling (Figure 2B). Although a cage effect was observed, it explained substantially less variation than diet (confirmed using PERMANOVA). Consistent with this, CD>WD mice clustered with long-term WD mice following the diet switch, whereas WD>CD mice clustered with long-term CD mice (Figure 2B-C). Longitudinal analysis showed no significant temporal changes in mice maintained on the same diet, whereas both diet-switch groups underwent significant microbial restructuring within 2 weeks of switching diets (PERMANOVA, p = 0.001; Figure 2C-D, Supplementary Figure 2L-M). This shift was more pronounced in the switch from chow to the western diet than from the western diet to chow diet (Figure 2C-D).

**Figure 2.**
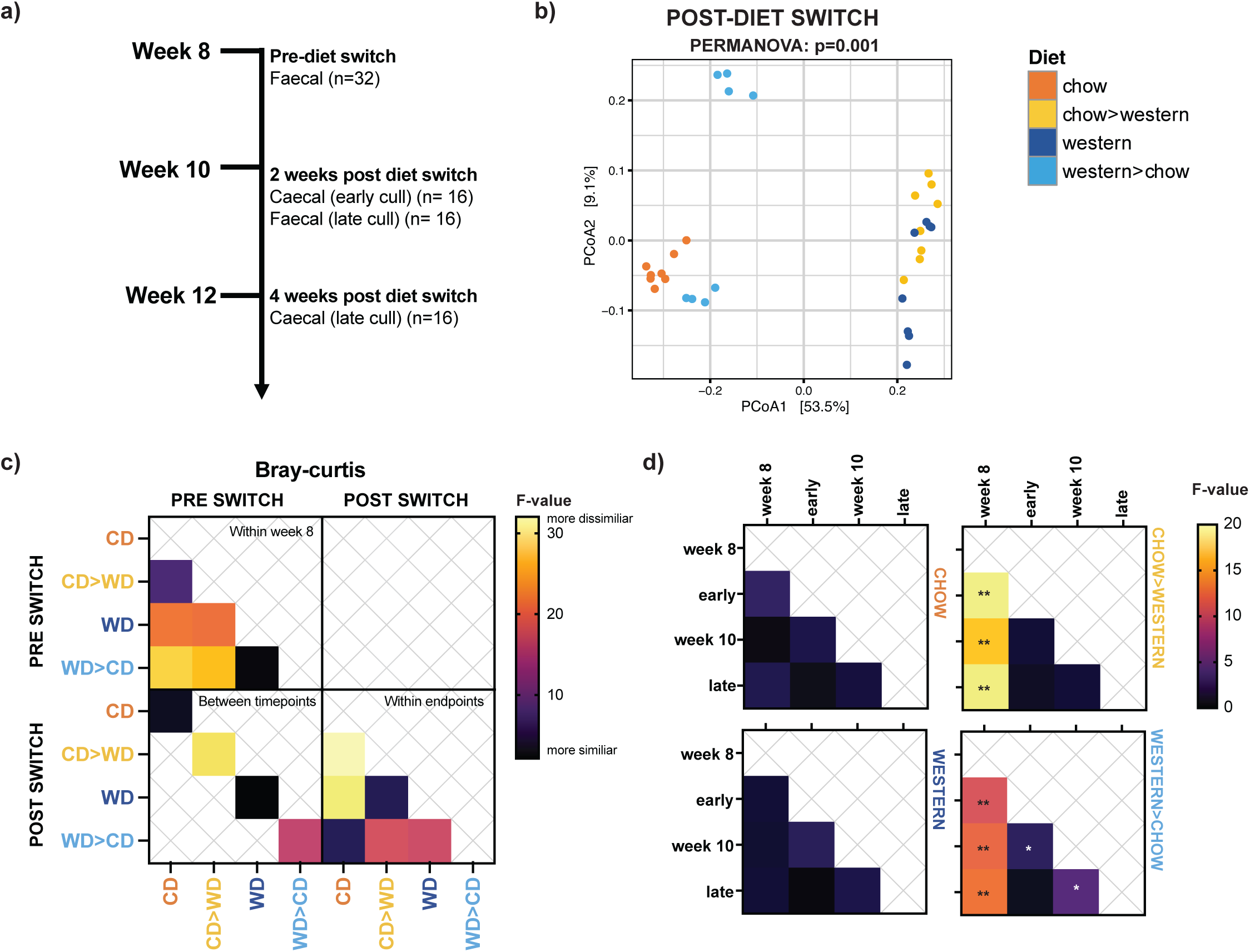
Diet switch rapidly remodels gut microbial community composition. (a) Overview of longitudinal microbiome sampling. (b) Principal coordinate analysis (PCoA) of beta diversity of microbiota samples after the diet switch (caecal samples week 10 or 12) calculated by Bray-Curtis dissimilarity. P-values were calculated using PERMANOVA. Coloured according to diet group: chow (orange), chow>western (yellow), western (dark blue) and western>chow (light blue). (c-d) Beta diversity across samples pre vs post diet switch were calculated by Bray-Curtis dissimilarity. Pairwise PERMANOVAs were conducted to determine statistical differences between all groups and timepoints, colours indicate F-values, p-values adjusted using the Benjamini-Hochberg method; * q< 0.05, ** q< 0.01, *** q<0.0001.

Pairwise differential abundance analysis (ANCOM-BC) using mice on long-term chow (CD) as the comparator demonstrated that the endpoint microbiota of CD>WD mice closely resembled long-term WD mice, whereas WD>CD mice exhibited substantial recovery of genera enriched in long-term CD-fed mice (Figure 3A). It was however evident that recovery was incomplete with in the short timeframe, with several taxa not recoverable to long-term CD-associated relative abundance including *Bifidobacterium*. Longitudinal analysis of genus-level abundance revealed largely reciprocal microbial changes in response to diet switching (CD>WD and WD>CD) and limited changes in mice that remained on CD or WD long term (Supplementary Figure 2A). The switch to the western diet was associated with a significant reduction in *Ligilactobacillus, Lachnospiraceae UCG-001*, *Muribaculum* and *Bifidobacterium* and an increase and enrichment in *Erysipelatoclostridium, Bilophila, Lactococcus lactis* and *Bacteroides thetaiotaomicron* (Figure 3A, Supplementary Figure 2A, 3A-J). In contrast the switch the chow diet was associated with a significant reduction in *Erysipelatoclostridium, Bilophila, Lactococcus lactis, Akkermansia muciniphila, Bacteroides thetaiotaomicron* and *Romboutsia ilealis* and a significant increase in *Ligilactobacillus*, *Lachnospiraceae UCG-001* and *Muribaculum* (Figure 3A, Supplementary Figure 2A, 3A-J).

**Figure 3.**
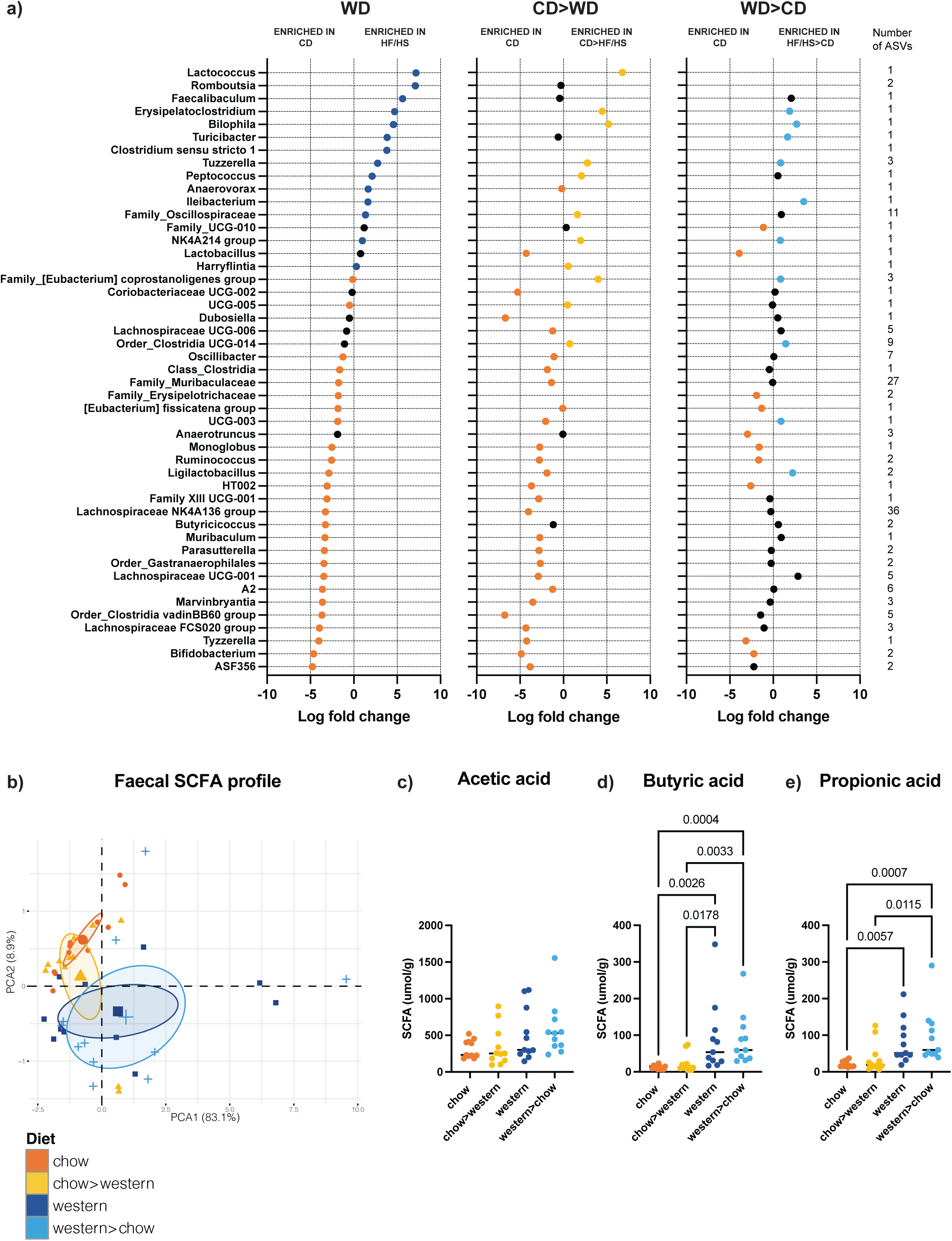
Gut microbial composition and faecal short-chain fatty acid profiles exhibit distinct responses to dietary switching. (a) Differential abundance of genus level taxa at endpoint assessed using ANCOM-BC between mice on the chow diet long term (chow)(n=8) and mice on that remained on the western diet (western)(n=8), mice that switched onto the western diet (chow>western) (n=8) or mice that switched onto the chow (western>chow) (n=8). Taxa which were significantly differentially abundant taxa in at least one group (q < 0.05) are presented as log2 fold change between mice that remained on the chow diet and the diet group. Dots are coloured if the taxa was significantly enriched (chow = orange, western= dark blue, chow>western = yellow, western>chow = light blue). Number of ASVs contributing to each genus are listed. (b) Principal component analysis (PCA) of overall quantified faecal SCFA profiles at endpoint, coloured by diet group. (c-e) Faecal (c) acetic acid, (d) butyrate and (e) propionate level assessed by LC-MS/MS between diet groups. Dots indicate individual mice, bars indicate median. P-values calculated by Kruskal-Wallis with post-hoc Dunn test for multiple comparisons

### Short-term dietary switching does not fully restore faecal SCFA profiles

To assess whether the rapid changes in microbial community composition were accompanied by changes in microbial metabolites, faecal SCFAs were quantified at endpoint by LC-MS/MS. Principal component analysis of the overall quantified SCFA profile demonstrated clustering primarily according to the diet the mice were on long-term, with mice switched to the alternate diet retaining SCFA profiles more similar to their original diet group than their current diet (Figure 3B). Consistent with this, faecal butyrate and propionate were significantly higher in WD-fed mice compared to CD mice and remained elevated following the switch from western to chow (WD>CD). Similarly, faecal butyrate and propionate levels in CD>WD mice remained lower than those observed in mice maintained on the western diet. Together, these findings indicate that unlike gut microbial composition, faecal SCFA profiles remained largely determined by long-term dietary exposure.

As the western diet was associated increased intestinal permeability prior to dietary switching, we next assessed whether intestinal barrier function altered following diet switching, 2 or 4 weeks later. A similar trend towards increased intestinal permeability with the Western diet compared to the chow was observed when mice were grouped according to the diet at the time of measurement (Supplementary Figure 2B), and when combining by timepoints (p = 0.0066; Supplementary Figure 2C). Although longitudinal measurements showed considerable inter-animal variability, mice switched from chow to the western diet exhibited a trend towards increased intestinal permeability (Supplementary Figure 2D-G). Collectively, these findings support an association between western diet consumption and impaired intestinal barrier function.

### Local and peripheral immune populations exhibit distinct temporal responses to dietary switching

To determine how the different diets influenced local and systemic immunity, single cell suspensions of immune cells were prepared from the mesenteric lymph nodes (MLN), blood and spleen at endpoint and comprehensively profiled using mass cytometry (35 marker panel) (Figure 4A). Unsupervised FlowSOM clustering was used to generate 40 metaclusters spanning T cell, B cell and myeloid populations (Figure 4B, Supplementary Figure 4A, Supplementary Table 1). Long term consumption of chow versus western diet established distinct immune landscapes across the MLN, PBMC and spleen (Supplementary Figure 4B-C). Differential abundance analysis identified significant differences in multiple immune metaclusters within the MLN and spleen, whereas comparatively few differences were observed in PBMCs, indicating that dietary effects were most pronounced within the gut-draining lymphoid compartment (Supplementary Figure 4D).

**Figure 4.**
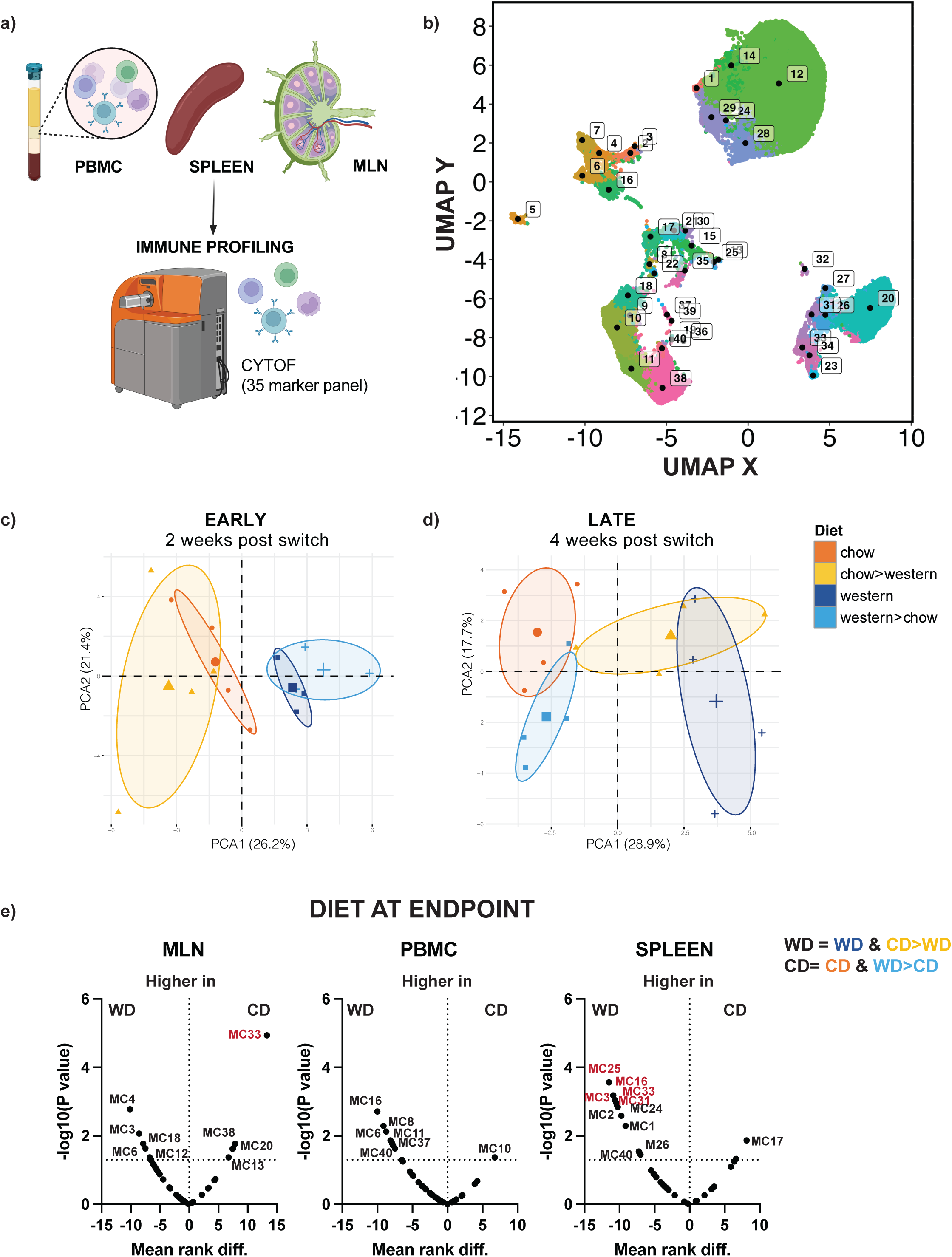
Dietary switching progressively remodels local and systemic immune landscapes. (a) Immune analysis overview. Immune cells were isolated from the MLN, blood and spleen of all mice at the experimental endpoint, stained with a panel of 35 markers and analysed using mass cytometry. Unsupervised FlowSOM clustering was used to generate 40 metaclusters spanning T cell, B cell and myeloid populations present across the MLN, PBMC and spleen of mice. (b) UMAP plot visualising major immune cell populations, coloured, and annotated according to metacluster. (c-d) Principal component analysis of overall immune repertoires at (c) 2 weeks and (d) 4 weeks post diet swap, coloured by diet group (chow = orange, western= dark blue, chow>western = yellow, western>chow = light blue). Volcano plots indicating metaclusters that are differentially represented between mice grouped according to the diet at the endpoint (chow & western>chow groups (n=16) vs western & chow>western groups (n=16)) across the MLN, PBMC and spleen. P-values calculated with Mann-Whitney U rank, p<0.05 are annotated. Metaclusters coloured in red pass post-hoc Dunn test for multiple comparisons.

We next investigated how the overall immune landscape changed following diet switching (Figure 4C-D). Principal component analysis revealed that 2 weeks after diet switching (CD>WD or WD>CD), the overall immune repertoires in the MLN of mice remained clustered according to the long-term priming diet. However, by 4 weeks post diet-switch samples clustered according to the diet at the endpoint, indicating progressive remodelling of the gut-draining immune compartment. We therefore examined individual immune populations by grouping mice according to either their long-term dietary exposure or the diet consumed at the experimental endpoint, allowing us to distinguish populations that rapidly remodelled following dietary switching from those that retained signatures of long-term dietary conditioning (Figure 4E, Supplementary Figure 4E).

We next examined major immune cell subsets using manual gating. Long term western diet feeding was associated with higher frequencies of overall B cells in both the MLN and spleen compared to mice on the chow diet long term (Supplementary Figure 5A). Following dietary switching, total B cell frequencies (% CD45+) remained associated with the long-term priming diet, with CD>WD mice resembling long-term chow-fed mice and WD>CD mice resembling long-term western diet-fed mice. In contrast, activation-associated B cell phenotypes rapidly remodelled following diet switching. Ki67+ B cells were significantly higher (% CD45) across the MLN, PBMCs and spleen of mice on the western diet at endpoint (p= 0.0067, p= 0.0084 and p <0.0001 respectively; Figure 5A-B). Similar changes were observed when expressed as a proportion of B cells within the PBMC and spleen (Supplementary Figure 5B). PD-1+ B cells exhibited a comparable pattern, with increased frequencies in the PBMCs and spleen of western diet-fed mice, both as a proportion of CD45+ cells and of total B cells (Figure 5C, Supplementary Figure 5C-D). Together, these findings indicate that B cell activation rapidly adapts to changes in dietary composition, in contrast to the persistence of total B cell abundance.

**Figure 5.**
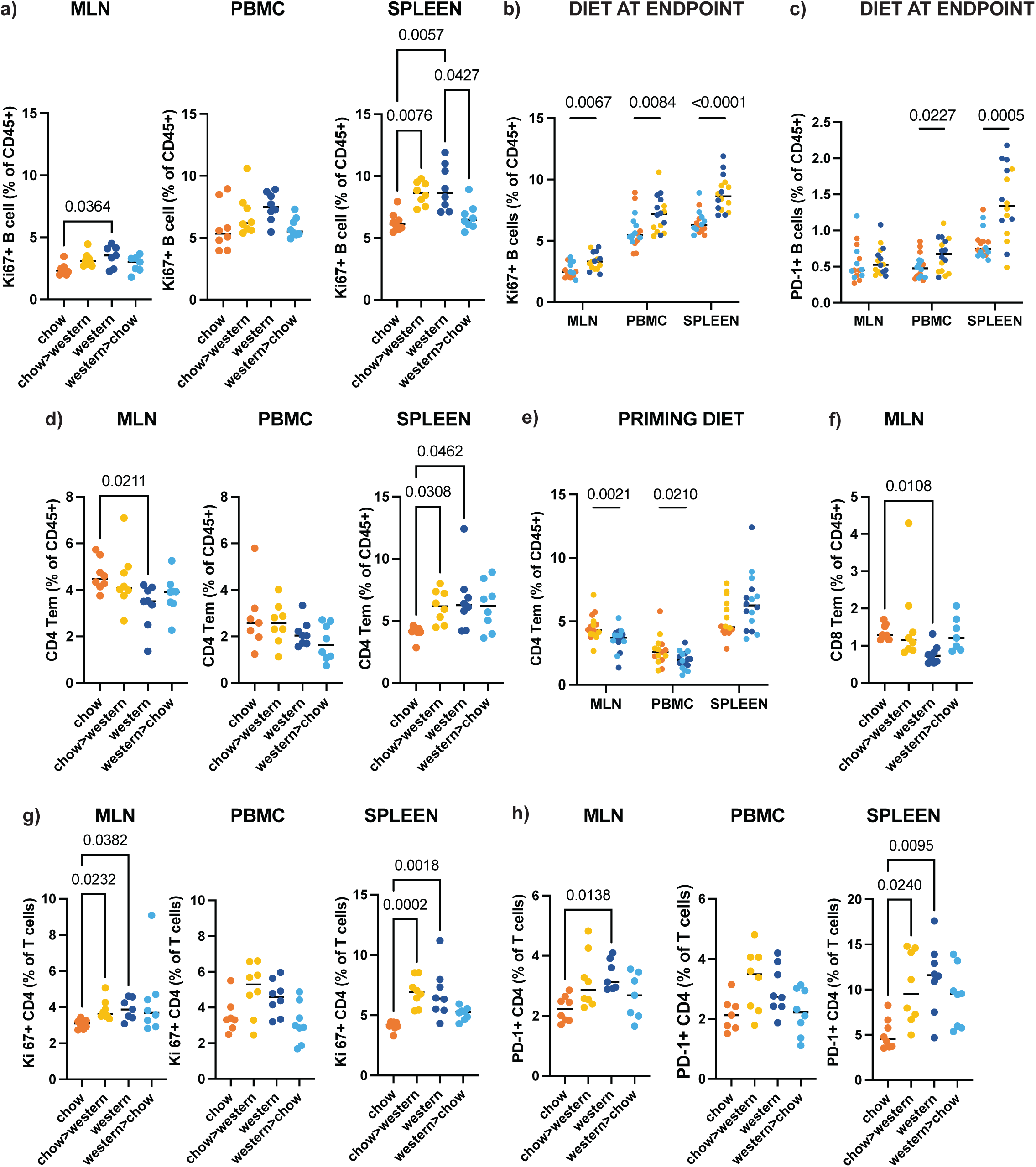
Activation-associated lymphocyte populations rapidly respond to dietary switching whereas memory T-cell populations retain signatures of long-term dietary exposure. (a-b) Frequency of Ki67+ B cells across MLN, PBMC or spleen (a) split per diet group (% CD45) or (b) pooled according to the diet at the endpoint (% CD45) (c) Frequency of PD-1+ B cells pooled according to the diet at the endpoint as a percentage of CD45+ cells (% CD45). (d-e) Frequency of CD4 + Tem (CD44+ CD62L) across MLN, PBMC or spleen (e) split per diet group or (e) pooled according to long term priming diet (% CD45). (f) Frequency of CD8+ Tem in MN split per diet group (% CD45). (g) Frequency of Ki67+ CD4 across MLN, PBMC or spleen split per diet group (% T cells). (h) Frequency of PD-1+ CD4 across MLN, PBMC spleen (% T cells). Coloured according to diet group: chow (orange), chow>western (yellow), western (dark blue) and western>chow (light blue). Dots indicate individual mice, bars indicate median. P-values calculated by Kruskal-Wallis with post-hoc Dunn test for multiple comparisons.

Similar dynamics were observed within the T cell compartment. CD4+ and CD8+ effector memory T (Tem; CD44+CD62L-) cells remained enriched in mice primed on the chow diet despite dietary switching, indicating that memory T cell populations largely retained signatures of long-term dietary exposure (Figure 5D-F, Supplementary Figure 5E-G). In contrast, activation-associated T cell phenotypes rapidly remodelled following diet swapping. Ki67+ CD4+ T cells and PD-1+ CD4 T cells were significantly increased in the MLN, PBMC and spleens of mice on the western diet at the endpoint (Figure 5G-H, Supplementary Figure H-I). Similar changes were observed within the CD8 T cell compartment, although these were primarily restricted to the MLN (Supplementary Figure 5J-M). Although PD-1+ CD8 T cells were enriched in the spleen only in mice maintained on the western diet long term (Supplementary Figure 5L). Collectively, these findings demonstrate that memory T cell populations retain a long-term dietary imprint, while activated CD4 and CD8 T cell subsets rapidly respond to changes in diet composition.

Regulatory immune populations also exhibited distinct temporal responses to diet switching. CD103+ dendritic cells (CD103+ cDC; MHCII+CD11c+) remained enriched in the MLN of mice primed on the chow diet following dietary switching (Figure 6A, Supplementary Figure 6A). Similarly, total Tregs (CD25+FoxP3+) were enriched in the MLN of mice primed on the chow diet (p=0.0049) (Figure 6B, Supplementary Figure 6B), whereas splenic Tregs were enriched in mice on the western diet at endpoint (p=0.0227) (Supplementary Figure 6C). In contrast, activated Treg subsets rapidly remodelled in response to short-term dietary switching. ICOS+ Treg subsets in particular revealed inverse associations with diet across sites, with ICOS+ Tregs being highly enriched in the MLN of mice on the chow diet (p< 0.0001) while ICOS+ Tregs were significantly enriched in the spleen of mice on the western diet at the endpoint (p= 0.0035) (Figure 6C, Supplementary Figure 6D). To further investigate Treg heterogeneity, unsupervised FlowSOM clustering analysis identified six Treg subtypes differing primarily in activation, proliferation and memory phenotypes (Supplementary Figure 7A-J). Manual gating confirmed that RORγt+ Tregs were rapidly restored following switching from the western to the chow diet, reaching frequencies comparable to long-term chow-fed mice within 2 weeks (Figure 6D, Supplementary Figure 7E-F). In contrast, resting Tregs were not altered by diet and ICOS+PD-1+Ki67+ Tregs were enriched in the spleen of mice on the western diet (Figure 6E, Supplementary Figure 7I-J).

**Figure 6.**
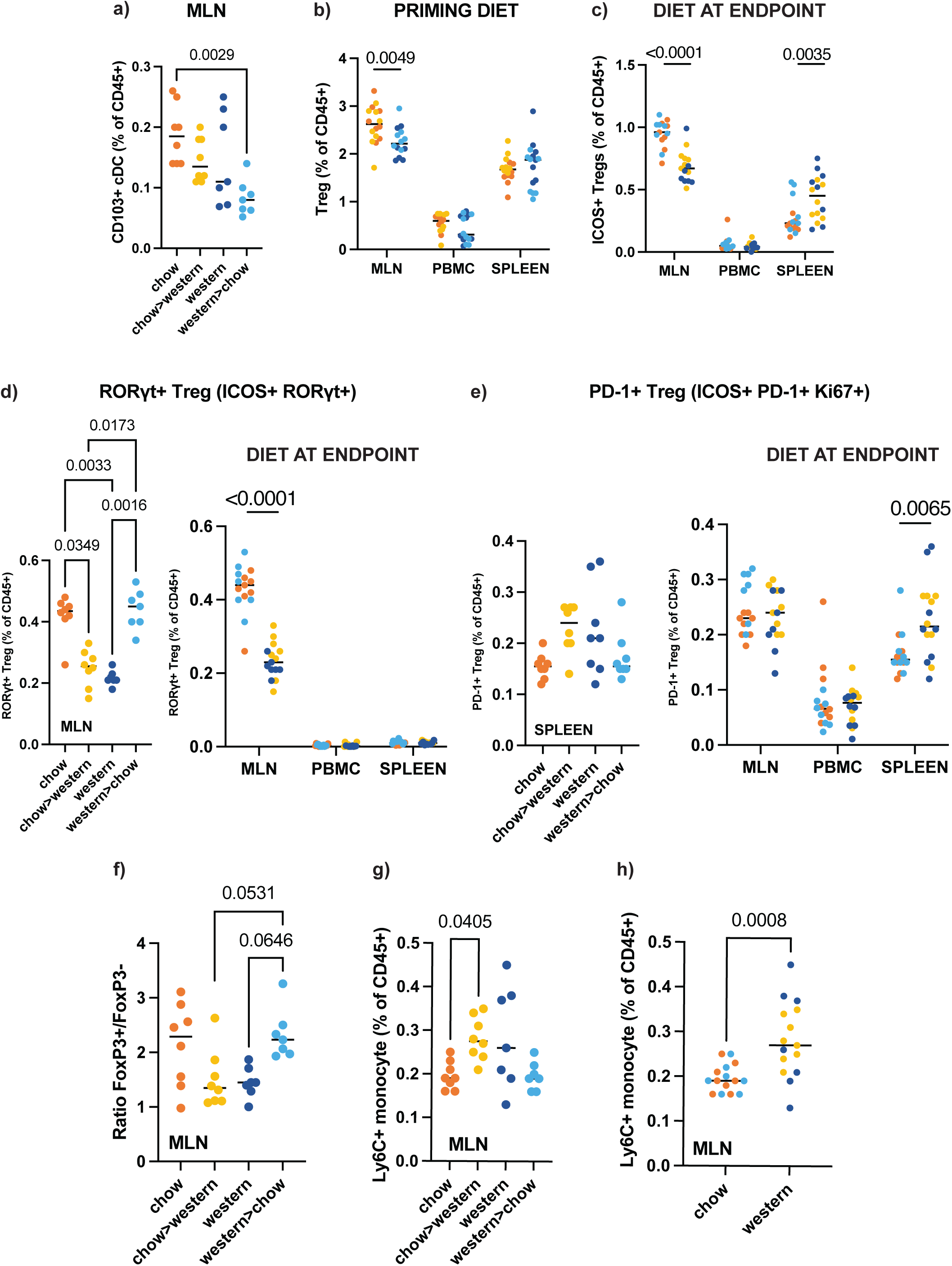
Dietary switching rapidly restores RORγt+ regulatory T cells whereas total regulatory populations retain signatures of long-term dietary exposure. (a) Frequency of CD103+ cDC (MHCII+ CD11c+) (% CD45) in the MLN split per diet group (b) Frequency of Tregs (CD25+ FoxP3+) (% CD45) across MLN, PBMC or spleen pooled according to long term priming diet. (c) Frequency of ICOS+ Tregs (% CD45) across MLN, PBMC or spleen pooled according to diet at the endpoint. (d) Frequency of RORγt+ Treg (ICOS+ RORγt+) across the MLN, PBMC or spleen, (left) split group or (right) pooled according to diet at the endpoint. (e) Frequency of PD-1+ Tregs (ICOS+ PD-1+ Ki67+) across the MLN, PBMC or spleen, (left) split group or (right) pooled according to diet at the endpoint. (f) Ratio of Treg/Th17. (g) Frequency of inflammatory monocytes (CD11b+Ly6C+) (% CD45) in the MLN. (f) Pooled frequency of inflammatory monocytes according to diet at the endpoint. Coloured according to diet group: chow (orange), chow>western (yellow), western (dark blue) and western>chow (light blue). Dots indicate individual mice, bars indicate median. P-values indicated, Mann-Whitney U rank or Kruskal-Wallis with post-hoc Dunn test for multiple comparisons presented where appropriate.

Given the rapid recovery of RORγt+ Tregs, we next examined the balance between regulatory and Th17 responses within the MLN. Although total RORγt expressing CD4 T cells were enriched in the MLN of mice on the chow diet at the experimental endpoint, there was no difference in conventional Th17 (RORγt+ FoxP3-) cell frequencies between diets (Supplementary Figure 6E-G). However, the proportion of RORγt+ CD4 expressing FoxP3 was greater in mice chow-fed mice (Figure 6F), suggesting a shift towards a more regulatory immune environment. To explore potential relationships between diet-associated microbial taxa and regulatory immune responses, correlations were performed between differentially abundant genera and MLN Treg subsets (Supplementary Figure 6H). Chow-associated genera including *Muribaculum, Ligilactobacillus* and members of the *Lachnospiraceae* family positively correlated with ICOS+RORγt+ Treg populations, whereas western diet-associated genera including *Erysipelatoclostridium, Tuzzerella* and *Bilophila* exhibited the opposite pattern and instead positively correlated with RORγt-Treg populations.

Finally, inflammatory monocytes within the MLN also rapidly responded to dietary switching, with higher frequencies observed in mice on the western diet at the experimental endpoint (Figure 6G-H). Collectively, these findings demonstrate that dietary switching differentially remodels immune populations, with activation-associated lymphocyte and inflammatory myeloid populations responding rapidly to changes in dietary composition, whereas several memory and regulatory immune populations retain signatures of long-term dietary exposure (Supplementary Table 2). These findings suggest that short-term dietary intervention can partially remodel immune function despite persistent immune imprinting established by long-term dietary exposure.

## Discussion

Despite numerous dietary intervention studies and trials across a variety of contexts, remarkably little is known about how rapidly established immune homeostasis can be remodelled by short-term dietary interevntion^20^. This study therefore aimed to assess the capacity of a short-term dietary switch following a long-term priming diet to alter established microbial community assemblages and reprogramme immune responses. We demonstrate that short-term dietary intervention rapidly remodels gut microbial community structure and immune phenotypes both locally in the gut and systemically. However, different components of the diet-microbiome-immune axis exhibited distinct response kinetics. Gut microbial community structure was rapidly remodelled within two weeks of the dietary switch, whereas microbial metabolic outputs, particularly faecal SCFA profiles, remained more strongly associated with long-term dietary exposure. Likewise, while many immune populations responded rapidly to the diet switch, including the restoration of RORγt+ Tregs in the MLN and reductions in hyperactivated B and T cell phenotypes, other populations, including overall B cell and Treg frequencies, effector memory T cells and CD103+ DC retained signatures of the priming diet. Together, these findings indicate that microbial community composition, metabolites and immune populations exhibit distinct temporal plasticity following dietary intervention. This has important implications for clinical settings in which dietary interventions must exert effects over relatively short timeframes, such as cancer immunotherapy^21^. Understanding which components of the diet-microbiota-immune axis can be rapidly remodelled, and which require more prolonged dietary intervention, will help inform the design and duration of future dietary strategies.

As expected, the chow and western diets resulted in compositionally distinct microbial profiles. The chow diet enriched fibre-degrading and SCFA-producing taxa^22–28^, whereas the western diet favoured bile-tolerant, mucin-degrading and alternative substrate-utilising microbes that are characteristic of low-fibre, high fat nutritional environment^29–34^. Diet switching rapidly remodelled these established microbial communities in both directions. Switching from western to chow resulted in a substantial recovery of the gut microbial community towards that of long-term chow-fed mice whereas switching from chow to western rapidly depleted many fibre-associated taxa. Consistent with the increased fibre availability, switching to chow from the western diet reduced the abundance of intestinal mucin degraders (*Akkermansia muciniphila* and *Bacteroides thetaiotaomicron*), while the opposite was observed for the switch to the western diet^29,30^. However, ecological recovery was incomplete. Taxa such as *Bifidobacterium* remained depleted following the switch to chow, suggesting that restoration depends not only on renewed substrate availability but also on the persistence of organisms capable of recolonising the community^35^. Conversely, an increase in key microbes enriched in western mice such as *Romboutsia* and *Faecalibaculum* was not observed in mice that switched from the chow to the western diet within 2-4 weeks, suggesting that prolonged dietary exposure, resulting in greater metabolic dysfunction or persistent protective effects of long-term chow priming may influence their expansion. Interestingly, a larger compositional shift was observed in mice that switched from CD to WD than from the WD to CD. This likely reflects the stronger ecological selection imposed by the western diet, whereby reduced nutrient availability rapidly depletes fibre-associated organisms and favours microbes adapted to alternative substrates such as host-derived mucins and bile acids. In contrast, recovery following the switch to chow may be constrained because key fibre-utilising taxa must first re-establish before they can fully exploit the increased availability of fermentable substrates

Despite rapid taxonomic restructuring, faecal SCFA profiles remained largely associated with long-term dietary exposure. Faecal butyrate and propionate concentrations remained higher in mice primed on the western diet, despite substantial recovery of fibre-associated microbial taxa following the switch to chow. This discordance indicates that rapid taxonomic remodelling is not necessarily accompanied by equivalent changes in the faecal metabolite pool. Although SCFA production is typically linked to dietary fibre fermentation, faecal SCFA concentrations reflect the net balance between microbial production, intestinal absorption and host utilisation^4,36^. Several mechanisms could explain this persistent elevation, including altered intestinal absorption, microbial utilisation or continued differences in microbial metabolic activity despite compositional recovery. Together, these observations indicate that microbial community composition, microbial metabolic output and immune remodelling occur on distinct temporal trajectories following dietary intervention.

Consistent with previous reports that western-style diets impair intestinal barrier and induce low level inflammation^37–39^, mice on the western diet long term exhibited greater intestinal permeability, together with reduced Treg frequencies in the MLN compared to mice on the chow diet, indicating altered barrier function and changes in local immune regulation. Although overall Treg frequencies did not recover following the switch to chow, they remained strongly associated with long-term dietary exposure, highest overall in mice that remained on chow and lowest in mice that remained on the western diet. Given colonic Tregs are regulated by SCFAs^19,40,41^ it is unsurprising that this population was enriched in the MLN of mice on the higher fibre chow diet enriched with SCFA-producing taxa. The combination of enhanced intestinal barrier function and increased Treg frequencies despite lower faecal SCFA concentrations is consistent with greater host uptake and utilisation of SCFAs, rather than reduced microbial production. Notably, RORγt+ Tregs exhibited remarkable plasticity, rapidly recovering following the switch to chow and declining following the switch to western diet. RORγt+ Tregs are specialised regulators of intestinal inflammation and immune homeostasis^42–44^. Their differentiation depends on microbial signals, with multiple bacterial species inducing RORγt expression and reduced frequencies observed in germ-free and antibiotic-treated mice^42,44^. The rapid recovery of RORγt+ Tregs despite persistent alterations in faecal SCFA profiles suggests that restoration of this regulatory population is not driven solely by SCFAs but may also involve other microbiota-derived metabolites or microbial-associated signals^42,44^.

Mice on the western diet had significantly higher proportions of B cells across the MLN and spleen, particularly Ki67+ B cells, suggesting enhanced B cell proliferation. Previous studies have reported variable effects of obesogenic diets on B cell abundance and function^45–48^, likely reflecting differences in dietary composition (fat vs sugar content) and metabolic state. In our study, increased B cell frequency and proliferation was also accompanied by elevated PD-1 expression, suggesting that expansion of the B cell compartment occurred alongside increased functional exhaustion. A similar pattern of enhanced activation, proliferation and exhaustion was also observed across the T cell compartment, consistent with previous reports linking western diet and diet-induced obesity with increased T cell exhaustion and dysfunction in both mice and humans^49–52^. Notably, Ki67 and PD-1 expressing cells across the B and T cell compartment rapidly responded to dietary switching, indicating that immune activation states and functional capacity of immune cells remain highly plastic and can be rapidly modified by diet.

In contrast to activation-associated immune phenotypes, several immune populations, including CD103+ DC, overall Tregs and CD4 and CD8 effector memory T cells, remained closely linked with the long-term priming diet despite dietary switching, consistent with persistent dietary immune imprinting. CD103+ DC remained enriched in the MLN of mice primed on the chow even with the switch to the western diet and were not restored following 2-4 weeks of chow feeding after long-term western diet exposure. CD103+ DC promote the differentiation of Tregs in the gut^53^ and have been demonstrated to be enhanced by high fibre diets via the gut microbiota and SCFAs^19^. In line with this observation, our mice that remained on the chow diet long term had the highest abundance of CD103+ DC and overall Tregs in the MLN. However, the recovery of these populations was not observed in mice that switched onto the chow diet within the timeframe of the experiment, suggesting that restoration of immune homeostasis following long-term western diet exposure occurs more slowly than remodelling of microbial community composition and activation-associated immune phenotypes. In contrast, priming on a higher fibre chow diet may offer a degree of enhanced protection associated with a more stable regulatory immune state. Previous studies have similarly demonstrated persistent innate immune imprinting following western diet exposure, whereby enhanced innate immune responses towards LPS remain elevated despite dietary normalisation^11^. Our findings extend this concept by suggesting that dietary immune imprinting is not limited to western diet-induced inflammation but may operate bidirectionally, with long-term chow feeding establishing regulatory immune states that are comparatively resistant to short-term dietary switches.

Correlation analyses further connected the microbiome and regulatory immune findings. Chow-associated genera, including *Muribaculum*, *Ligilactobacillus* and members of the *Lachnospiraceae*, positively correlated with ICOS+RORγt+ Treg populations, whereas western diet-associated taxa such as *Bilophila*, *Erysipelatoclostridium* and *Tuzzerella* showed the opposite pattern. These associations were strongest in the MLN, consistent with its anatomical relationship to the intestine. However, both microbial taxa and immune populations were strongly structured by diet, and the correlations may therefore partly reflect their shared response to the intervention rather than direct microbe-immune interactions. Candidate organisms emerging from these analyses will require testing through defined microbial supplementation or transfer experiments.

This study has several limitations. The chow and western diets differed in multiple dietary components, including sugar, fat and fibre content and source, preventing attribution of the observed effects to any single macronutrient. Furthermore, the diets were not isocaloric and mice on the western diet exhibited greater weight gain, indicating that alterations in host metabolism associated with diet-induced obesity may have contributed to the observed immune phenotypes^15,25,54^. However, immune remodelling was evident within two weeks of diet switching, suggesting that a least some of these changes occur before the development of more prolonged metabolic dysfunction. The relative contributions of dietary nutrients, host metabolism and microbial restructuring cannot be resolved without complementary approaches, such as isocaloric diets, microbiota transfer, antibiotic treatment or gnotobiotic models. Nevertheless, extensive experimental evidence indicates that the gut microbiota is a key mediator of diet-induced changes in intestinal barrier function and immune homeostasis ^25,37,38,55,56^. Finally, the study used adult male SPF mice that were weaned onto chow. Consequently, the degree of microbial and immune plasticity observed here may differ following early-life or multigenerational western diet exposure, in female mice, or under inflammatory or disease conditions.

Overall, this study demonstrates that short-term dietary intervention rapidly remodels the gut microbiota and selectively reprogrammes immune phenotypes, but complete restoration of immune homeostasis occurs more slowly than changes in microbial composition. Rather than responding as a single interconnected system, the diet-microbiome-immune axis exhibits distinct temporal trajectories, with microbial community composition, microbial metabolic output and immune populations displaying different degrees of plasticity following dietary intervention. These findings provide a framework for understanding how dietary interventions reshape host-microbiome interactions and suggest that although short-term dietary modification can rapidly improve aspects of immune regulation, sustained dietary change is likely required to fully reverse long-term dietary immune imprinting. Collectively, these findings highlight the therapeutic potential of dietary interventions to modulate host-microbiome interactions and immune homeostasis across a broad range of disease settings.

## Methods

### Animals & diets

All animal experiments were approved by the University of Sydney Animal Ethics Committee (AEC2020/1811 and AEC2022/2161) and performed in accordance with institutional guidelines. Male C57BL/6 mice (8 weeks old) were obtained from Australian BioResources (NSW, Australia) and housed under specific pathogen-free conditions at the Charles Perkins Centre, The University of Sydney (4 mice per cage). Mice were fed either a standard chow (SF00-100) or a Western diet (High Fat/High Sugar) (SF03-020) ad libitum purchased from Specialty Feeds, Glen Forest, Australia. Following eight weeks on their allocated diet, mice either remained on the same diet or switched to the alternate diet for two or four weeks. Diet transitions were performed over three days to minimise dietary stress. Body weight and food intake were monitored throughout the study.

### FITC-Dextran Intestinal Permeability Assay

Intestinal permeability was assessed longitudinally using the FITC-Dextran intestinal permeability assay. Following a 4 hour fast, mice received 500 mg/kg 4.4 kDa FITC-Dextran (Sigma-Aldrich) via oral gavage. Serum was collected 4 hours later and stored at-80°C in the dark until the assay was performed (once all longitudinal samples were collected). Serum from non-gavaged mice was diluted in PBS (1:4) and an 11-point FITC-Dextran standard curve was prepared via serial dilution (from 1250 μg/mL). Fluorescence was quantified from all samples and the standards using a TECAN fluorescence plate reader (excitation 450 nm and emission 520nm).

### Faecal DNA extraction and 16S rRNA gene sequencing

Genomic DNA was extracted from faecal or caecal contents samples from both mice using the FastDNA Spin Kit for Feces (MP Biomedical) as per the manufacturers protocol. Extracted DNA was stored at-80°C. DNA concentration was quantified using Qubit BR dsDNA assay kit (Invitrogen) and samples were diluted to 10 ng/μL. 16S rRNA gene amplicon sequencing was performed on all DNA samples. Barcoded amplicon libraries spanning the V4 hypervariable region of the 16S rRNA gene were prepared (515F-806R primer set-515F: GTGYCAGCMGCCGCGGTAA, 806R: GGACTACNVGGGTWTCTAAT) and sequenced using the Illumina MiSeq v2 2 x 250 bp platform at the Ramacotti Centre for Genomics (UNSW, Sydney, Australia). Sequencing data was analysed using R. Raw reads were processed using DADA2 which uses error profiles to define amplicon sequence variants (ASVs)^57^. Taxonomy was assigned to ASVs using the SILVA (v138.1) reference database. Prior to downstream analysis, ASVs that were less than 0.01% of total reads and present in fewer than 5% of samples were filtered from the final dataset. Sequencing depth analyses and rarefaction were performed with the *phyloseq* R package^58^. Analysis and presentation of the resultant ASVs were performed in R using the packages *phyloseq*, *vegan*, *microbiome* and *ggplot2*. Alpha diversity metrics were calculated using Inverse Simpson’s index, species richness and Shannon’s index. Beta diversity was assessed on centred-log-ratio transformed ASV counts using Bray Curtis dissimilarity and UniFrac distance and principle coordinate analysis (PCoA) performed and plotted for visualisation. PERMANOVA (adonis) using the *vegan* R package was used to assess variance in the distance matrices between groups (i.e. the degree to which samples cluster based on microbial profile). Differential abundance analysis was also performed using the *ANCOM-BC* R package^59^.

### Preparation of single cell suspensions

Single cell suspensions were prepared from spleen and MLN from mice for immune analysis. Spleen and MLN were finely chopped and incubated with Collagenase Type IV and DNase (ThermoFisher) (1 mg/mL and 1U respectively in RPMI (10% FCS)). Digested samples were gently pressed through a 70uM strainer to produce single cell suspensions. ACK lysis buffer was used to lyse red blood cells in the spleen. PBMCs were isolated from blood collected from mice at endpoint using Histopaque 1083 (Sigma-Aldrich) density gradient.

### Mass cytometry (CyTOF)

For immune profiling of mouse MLN, PBMC and spleen, samples were stained with a panel of 35 antibodies (Table 1). Antibody cocktails were made up at pre-optimised concentrations in FACS (surface stains) or Perm/Wash buffer (intracellular) by Sydney Cytometry, The University of Sydney. For staining, cells were seeded into a round bottom 96 well plate (approximately 2 million per sample). Cell were centrifuged at 1500 rpm for 5 minutes and resuspended in 20 mg/mL purified anti-mouse CD16/32 antibody (Biolegend) and incubated on ice for 15 minutes. Cells were then washed with FACS and resuspended in 100 μL of barcoding antibodies (CD45-104, CD45-106 and CD45-108) and incubated for 20 minutes. CD45 antibodies labelled with 3 different metals (104, 106, 108) were used to barcode and combine cells obtained from MLN (CD45-104), PBMC (CD45-106) and spleen (CD45-108) from an individual mouse. Cells were then centrifuged and resuspended in Cell-ID Cisplatin (1:1000 in RPMI) (Fluidigm) and incubated for 3 minutes to stain dead cells, followed by quenching with FCS. Cells were then wash and resuspended in the surface antibody cocktail made up in 100 μL of FACS per sample (Table 1) and incubated on ice for 20 minutes. To prepare cells for intracellular staining, samples were resuspended in Fix/Perm buffer (1:4 fix/perm buffer to FoxP3 fixation/permeabilisation diluent) (Invitrogen, #00-5523-00) and incubated for 40 minutes. Cells were resuspended in the intracellular antibody cocktail made up in 100 μL of Perm/Wash buffer per sample (Table 1). Cells were then washed twice with Perm/Wash buffer and once with FACS. To distinguish cells from beads during sample acquisition, samples were resuspended in DNA intercalator (Cell-ID Intercalator-Ir, 1:4000 in 4% PFA) (Fluidigm) and incubated at 4°C overnight. Priro to acquisition cells were washed once with milliQ and once with CAS buffer (Cell Acquisition Solution for CyTOF) (Fluidigm). Cells were resuspended in CAS buffer and EQ Four Element Calibration Beads (Fluidigm) at 11% of the total volume. Samples were run on a Helios mass cytometer at Sydney Cytometry, The University of Sydney.

**Table 1.** Mouse CyTOF Panel.

| Antigen | Clone | Label | Stain |
| --- | --- | --- | --- |
| CD24 | 30-F1 | 89 | surface |
| CD45 | 30-F11 | 104 | surface |
| CD45 | 30-F11 | 106 | surface |
| CD45 | 30-F11 | 108 | surface |
| CD38 | 90 | 115 | surface |
| Ly6G | 1A8 | 141 | surface |
| CD11c | N418 | 142 | surface |
| CD194 (CCR4) | 2G12 | 143 | surface |
| CD4 | RM4-5 | 145 | surface |
| CD27 | LG.3A10 | 146 | surface |
| CD103 | 2E7 | 147 | surface |
| CD11b | M1/70 | 148 | surface |
| CD19 | 6D5 | 149 | surface |
| a4b7 | FIB504 | 150 | surface |
| CD25 | 3C7 | 151 | surface |
| CD3e | 145-2C11 | 152 | surface |
| CD278 (ICOS) | 7E.17G9 | 154 | surface |
| CD199 (CCR9) | 9B1 | 155 | surface |
| CD183 (CXCR3) | CXCR3-173 | 156 | surface |
| FoxP3 | FJK-16s | 158 | intracellular |
| CD45R/B220 | RA3-6B2 | 159 | surface |
| CD62L | MEL-14 | 160 | surface |
| CD279 (PD-1) | 29F.1A12 | 161 | surface |
| Ki67 | 11F6 | 162 | intracellular |
| CD44 | IM7 | 163 | surface |
| GATA3 | L50-823 | 164 | intracellular |
| CD196 (CCR6) | 29-2L17 | 166 | surface |
| CD185 (CXCR5) | 2GB | 167 | surface |
| CD8a | 53-6.7 | 168 | surface |
| TCR$\gamma\delta$ | GL3 | 169 | surface |
| NK1.1 | PK136 | 170 | surface |
| GZMB | REA226 | 171 | intracellular |
| RORγT | Q31-378 | 172 | intracellular |
| CD127 | A7R34 | 173 | surface |
| I-A/I-E (MHC-II) | M5/114.15.2 | 174 | surface |
| TIM-3 | RMT3-23 | 175 | surface |
| Ly6C | HK1.4 | 176 | surface |
| Tbet | 4B10 | 209 | intracellular |

### Cytometry Data Analysis

All FCS files were exported, and pre-gated using FlowJo (v10.8.1). Beads were first excluded followed by defining intact (DNA +ve) and live cells (cisplatin-ve) before debarcoding using CD45-104/106/108. Manual gating was conducted using FlowJo. The live leukocytes (CD45+ cells) datasets were also exported for downstream computational analysis using *Spectre* in R^60^. FlowSOM clustering and dimensionality reduction analysis was then performed and uniform manifold approximation and projection (U-MAP) plots were generated^61^. These were generated after FlowSOM clustering using down sampled data of 70,000 cells. FlowSOM clustering was performed using all markers. All significant clusters were confirmed using manual gating in FlowJo. Pairwise multiple Mann-Whitney tests were performed to determine differentially represented clusters or cell types between groups.

### SCFA derivatization and analysis

SCFA derivatization and analysis was performed as outlined in McKay *et al* 2023^62^. Faecal samples from mice were weighed and milliQ water was added to a final stool concentration of 5% w/v. Samples were then vortexed for 5 minutes until pellet was dissociated and centrifuged for 15 minutes at 14000 g. Faecal supernatant containing metabolites was transferred to a fresh tube and the remaining stool pellet was freeze dried overnight to determine the dry weight of the faecal sample. 10 μL of the faecal supernatant was used for derivatization. Samples were derivatized to 3-phenoxyaniline (3-PA) using 25 μL of 0.1M EDC in methanol and 10 μL of 0.5M 3-PA in methanol. Samples were then shaken for 30 minutes at room temperature to allow for carboxyl-to-amine coupling. 5 μL of trifluoroacetic acid was then added to quench the reaction and SCFA derivatives were purified by partitioning with 1 mL of tert-methyl ether. The ether layer was then removed and dried overnight, and samples were resuspended in 1:1 acetonitrile: water before analysis was conducted. LC-MS/MS was performed using the Shimadzu Nexera X2 UHPLC system.

## Acknowledgements

We acknowledge the technical assistance from Sydney Cytometry for mass cytometry, Laboratory Animal Services at the Charles Perkins Centre for animal housing and the Ramacotti Centre for Genomics for 16S sequencing. This work was supported by a Charles Perkins Centre EMCR Seed Funding grant and an NHMRC Ideas grant. R.C.S acknowledges financial support from Cancer Institute NSW. E.R.S. acknowledges financial support from the The William Arthur Martin à Beckett Cancer Research Trust (University of Sydney Fellowship).

## Supplementary Tables and Figures

**Supplementary Table 1.** Phenotypes of metaclusters with significant associations between diet groups.

| Metacluster | Phenotypic definition | Cell type |
| --- | --- | --- |
| MC1 | Ki67+ PD-1+ GrzB+ B220+ MHCII+ | PD-1+ B cell |
| MC2 | CD11c+ NK1.1+ IFN-γ+ MHCII+ CD16+ CD11b+ | NK/DC |
| MC3 | CD11c+ CD8+ NK1.1+ IFN-γ+ MHCII+ CD16+ | cDC1 |
| MC4 | CD11b+ Ly6C+ MHCII+ Ki67+ | Inflammatory monocyte |
| MC6 | CD11b+ Ly6C+ a4b7+ IFN-γ+ Ki67+CD16+ CD24+ | Gut homing inflammatory monocyte |
| MC8 | CD11c+ Ly6C+ Ki67+ | Ly6C+ DC |
| MC10 | CD3+ CD8+ CD27+ CD44+ CD62L+ Ly6C+ | CD8+ Tcm |
| MC11 | CD3+ CD8+ CD27+ CD62L+ Ly6C+ | Naïve CD8 T cell |
| MC12 | B220+ MHCII+ CD38+ CD44+ CCR6+ | Activated B cell |
| MC13 | CD3- MHCII+ RORγt+ CD25+ Granzyme B+ CD127+ CCR6+ | ILC3 |
| MC14 | B220+ MHCII+ CD38+CXCR5+ | Follicular B cell |
| MC16 | CD11b+ CD16+ CD44+ CD11c+ | cDC2 |
| MC20 | CD3+ CD4+ CD44+ CD62L+ | CD4+ Tcm |
| MC22 | CD11c+ B220+ CD38+ Ly6C+ IFNg+CCR9+ | pDC |
| MC24 | B220+ MHCII+ CD16+ CD38+ Ly6C+ | Plasma B cell |
| MC25 | CD3+ CD4- CD8- CD127+ RORγt+ CD44+ | MAIT |
| MC26 | CD3+ CD4+ CD44+ CD62L- PD-1+ CD27+ | PD-1+ CD4 Tem |
| MC27 | CD3+ CD4+ CD44+ CD62L- | CD4+ Tem |
| MC31 | CD3+ CD4+ CD44+ CD62L- PD-1+ Ki67+ | Proliferating PD-1+ CD4+ Tem |
| MC33 | CD3+ CD25+ FoxP3+ CD103+ ICOS+ CD127+ CCR6+ PD-1+ | ICOS+ Treg |
| MC34 | CD3+ CD25+ FoxP3+ ICOS- | ICOS- Treg |
| MC38 | CD3+ CD8+ CD62L+ CD103+ CD127+ CD27+ | CD8+ Trm |
| MC40 | CD3+ CD8+ CD44+CD62L+ Ki67+ PD-1+ | Proliferating CD8+ Tcm |

**Supplementary Table 2.**

| Site | Cell populations different between long term chow vs western |  | Cell populations responsive to diet changes |  |
| --- | --- | --- | --- | --- |
| **MLN** | **CD:**<br>↑ cDC (MHCII+ CD11c+)<br>↑ CD103+ cDC<br>↑ CD4 Tem<br>↑ CD8 Tem<br>↑ Tregs<br>↑ RORγt+ ICOS+ Tregs | **WD:**<br>↑ B cells<br>↑ Ki67+B cells<br>↑ Ki67+ CD4 T cells<br>↑ PD-1+ CD4 T cells<br>↑ Ki67+ CD8 T cells<br>↑ PD-1+ CD8 T cells | **CD:**<br>↑ RORγt+ Treg | **WD:**<br>↑ inflammatory monocytes (CD11b+ Ly6C+)<br>↑ Ki67+B cells<br>↑ Ki67+ CD4 T cells<br>↑ PD-1+ CD4 T cells<br>↑ Ki67+ CD8 T cells<br>↑ PD-1+ CD8 T cells |
| **PBMC** |  |  |  | **WD:**<br>↑ Ki67+B cells<br>↑ PD-1+ B cells<br>↑ Ki67+ CD4 T cells<br>↑ PD-1+ CD4 T cells |
| **SPLEEN** |  | **WD:**<br>↑ B cells<br>↑ Ki67+B cells<br>↑ PD-1+ B cells<br>↑ Ki67+ CD4 T cells<br>↑ PD-1+ CD4 T cells<br>↑ PD-1+ CD8 T cells<br>↑ CD4 Tem<br>↑ Tregs |  | **WD:**<br>↑ Ki67+B cells<br>↑ PD-1+ B cells<br>↑ Ki67+ CD4 T cells<br>↑ PD-1+ CD4 T cells<br>↑ PD-1+ Tregs |

---

**Supplementary Figure 1.**
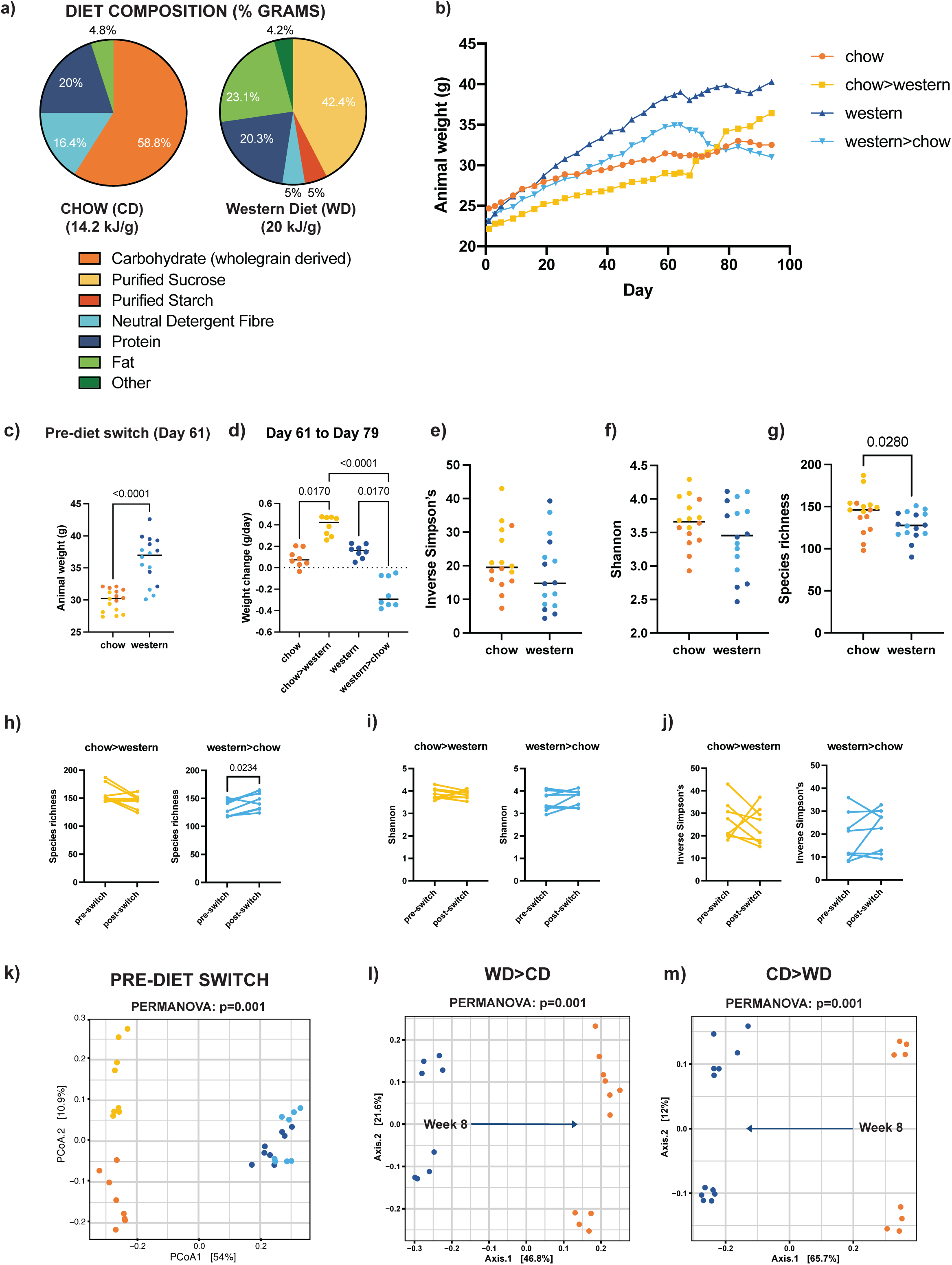
(a)Diet composition (% grams) of the chow (CD) and Western diets (WD). (b) Average body weight per group over the experiment. (c) Animal weight per mouse pre-diet switch (Day 61), grouped according to long-term diet, coloured according to the diet group (switched/ remained on). (d) Average weight change per day in the 2 weeks post diet switch (Day 61 to Day 79). (e-f) Alpha diversity calculated by (e) Inverse Simpson’s, (f)Shannon Indexes and (g) Species richness with samples grouped by long-term diet and coloured by diet group (switched onto or remain on) (orange= CD, yellow= CD>WD, dark blue= WD, light blue= WD>CD). Dots indicate individual mice, bars indicate median. P-values calculated by Mann-Whitney U rank test. (h-j) Change in alpha diversity between pre-diet switch faecal samples (week 8) and caecal samples collected at the respective endpoints (2 or 4 weeks later) of mice in the diet switch groups calculated by (h) species richness, (i) Inverse Simpson’s and (j) Shannon Indexes. Chow to western (yellow), Western to chow (light blue). Dots indicate individual mice, p-values of paired samples assessed by Wilcoxon sign-ranked test. (k) Principal coordinate analysis (PCoA) of beta diversity of microbiome samples calculated by Bray-Curtis dissimilarity. Coloured according to diet group (orange= CD, yellow= CD>WD, dark blue= WD, light blue= WD>CD).(l-m) Principal coordinate analysis (PCoA) of beta diversity of microbiota samples before and after the diet switch calculated by Bray-Curtis dissimilarity in (j) WD>CD and (k) CD>WD mice. P-values were calculated using PERMANOVA.

**Supplementary Figure 2.**
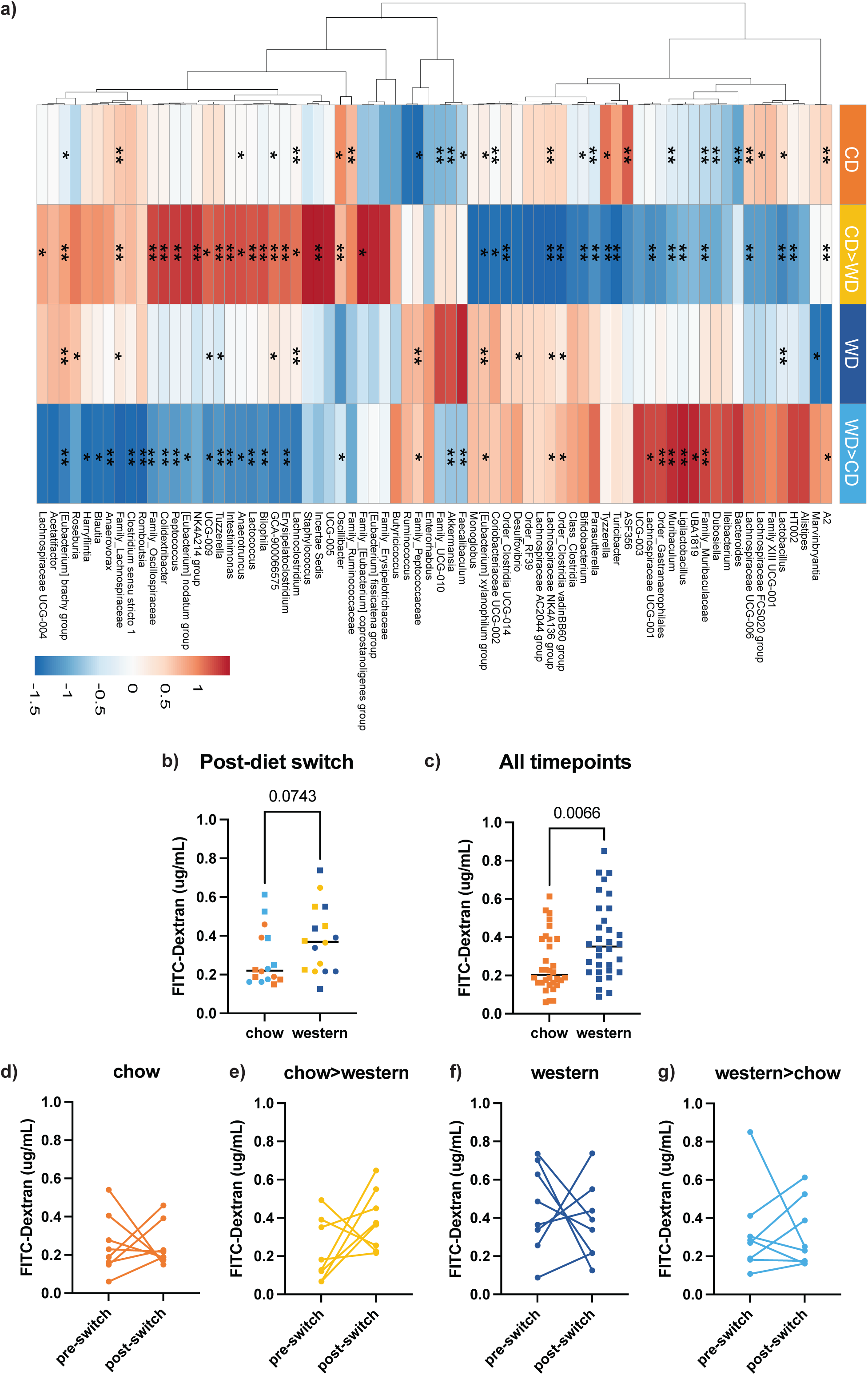
(a) Log fold change relative abundance of genus level taxa between pre-diet switch faecal samples and post diet switch caecal samples was assessed (log2, caecal/week 8). Heatmap presents average fold change per group, row centred to indicate extent of change between timepoints. Wilcoxon sign-ranked test performed on individual paired samples within each diet group, asterisks indicate p-values, * p< 0.05, ** p< 0.01 (Benjamini-Hochberg corrected, all q<0.1). (b) Post-diet switch FITC-Dextran in serum samples of mice is indicated, with datapoints grouped according to the diet at the time of the sampling. Colours indicate the diet group the mice belong to (circle= week 10, square= week 12). (c) Pooled data from all timepoints grouped according to the diet at the time of sampling. (d-g) Longitudinal change in intestinal permeability between pre-diet switch timepoint and endpoint (post-diet switch) per mouse within each diet group. Dots indicate individual mice, bars indicate median. P-values indicated (only values p < 0.1 shown), calculated by Mann-Whitney U rank test or Wilcoxon sign-ranked test where appropriate.

**Supplementary Figure 3.**
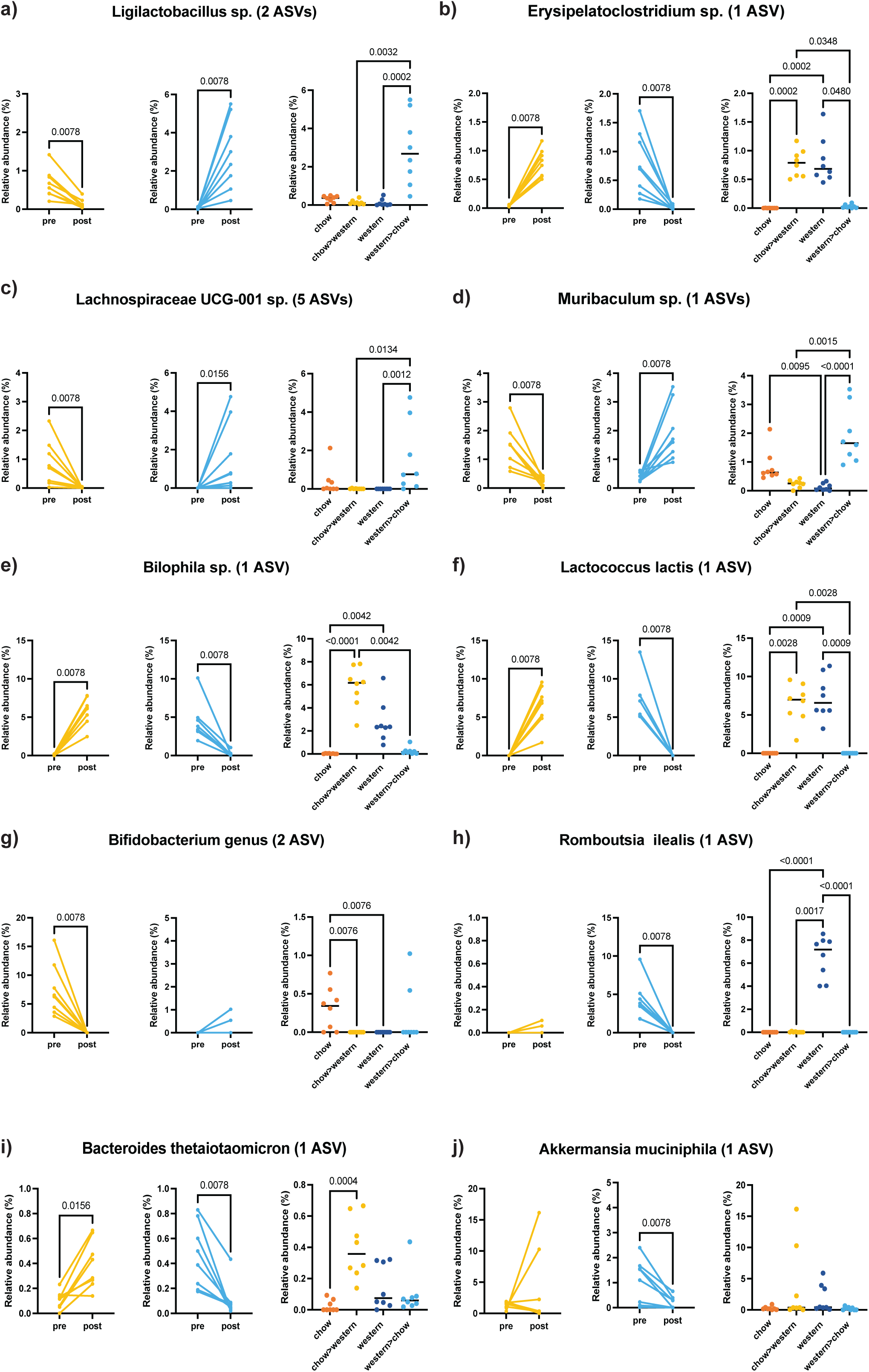
(a-j) Selected taxa that exemplify different patterns of change in relative abundance in the diet switch groups (chow>western in yellow, western>chow in light blue) and comparative relative abundance of taxa at the endpoint across all 4 diet groups (chow = orange, western= dark blue, chow>western = yellow, western>chow = light blue). Dots indicate individual mice, bars indicate median. P-values are indicated. Wilcoxon sign-ranked test or Kruskal-Wallis with post-hoc Dunn test for multiple comparisons where appropriate.

**Supplementary Figure 4.**
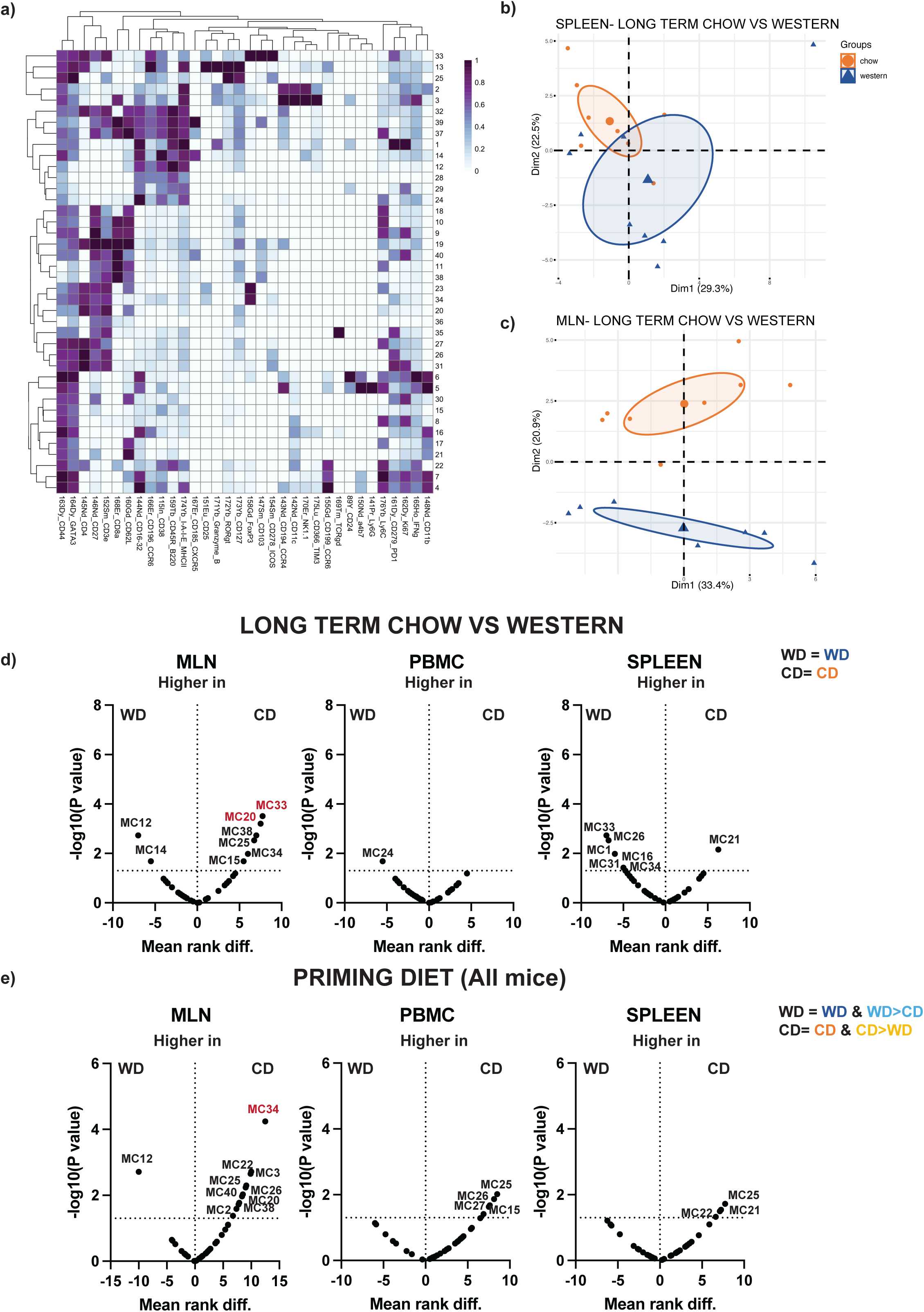
(a) Heatmap showing relative median signal intensity of markers for each metacluster from FlowSOM unsupervised clustering analysis. (b-c) Principal component analysis of overall immune repertoires in the (b) spleen or (c) MLN of mice that remained on the chow or western diets long term. (d) Volcano plots indicating metaclusters that are differentially represented between mice that remained on the chow diet long term (chow n=8) and mice that remained on the western diet long term (western n=8) across the MLN, PBMC and spleen (e) Volcano plots indicating metaclusters that are differentially represented between mice that were primed on the chow diet and mice primed on the western diet (long term diet phase) (chow & chow>western groups (n=16) vs western & western>chow groups (n=16)) across the MLN, PBMC and spleen. P-values calculated with Mann-Whitney U rank, p<0.05 are annotated. Metaclusters coloured in red pass post-hoc Dunn test for multiple comparisons.

**Supplementary Figure 5.**
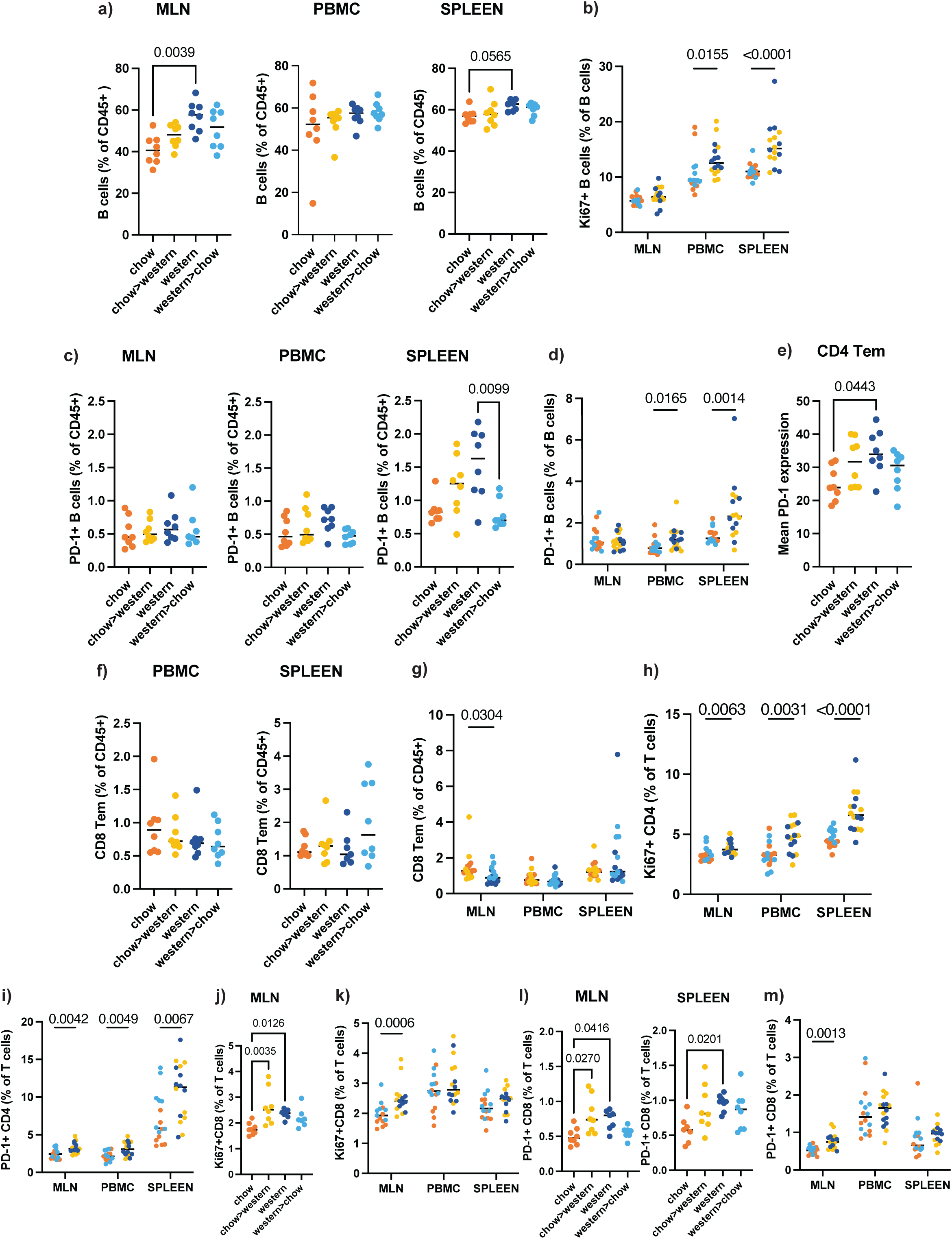
(a) Frequency of B cells (MHCII+ B220+) (%CD45) across MLN, PBMC or spleen. (b) Frequency of Ki67+ B cells across MLN, PBMC or spleen pooled according to the diet at the endpoint as percentage of B cells. (c-d) Frequency of PD-1+ B cells across MLN, PBMC or spleen (c) split per diet group (%CD45) or (e) pooled according to the diet at the endpoint (% B cells). (e) Mean PD-1 expression on CD4+ Tem in the spleen grouped according to diet group. (f-g) Frequency of CD8+ Tem (CD44+ CD62L-) as a percentage of CD45+ cell across PBMC or spleen (f) split per diet group or (g) pooled according to long term priming diet. (h) Frequency of Ki67+ CD4 T cells across MLN, PBMC or spleen pooled according to diet at the endpoint. (% T cells). (i) Frequency of PD-1+ CD4 T cells across MLN, PBMC or spleen pooled according to diet at the endpoint (% T cells). (j) Frequency of Ki67+ CD8 T cells (% T cells) in the MLN split per diet group or (k) pooled according to diet at the endpoint. (l-m) Frequency of PD-1+ CD8 T cells across MLN, PBMC or spleen (l) split by diet group or (m) pooled according to diet at the endpoint (% T cells). Dots indicate individual mice, bars indicate median. P-values are indicated. Kruskal-Wallis with post-hoc Dunn test for multiple comparisons.

**Supplementary Figure 6.**
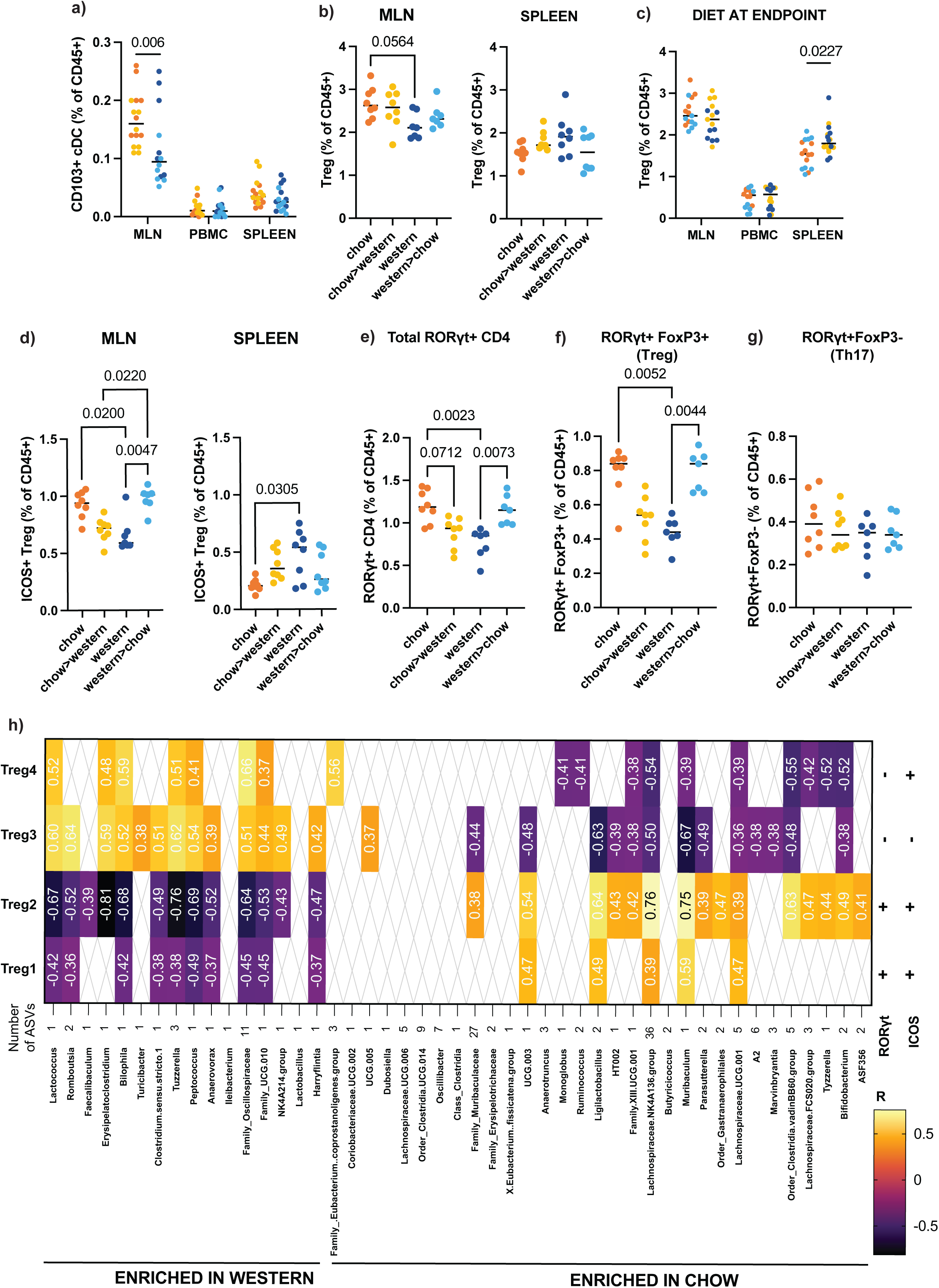
(a) Frequency of CD103+ cDC (% CD45) across the MLN, PBMC, spleen group according to the long-term priming diet. (b-c) Frequency of Tregs (CD25+ FoxP3+) (%CD45) across MLN, PBMC or spleen (b) split per diet group or (c) pooled according to diet at the endpoint. (d) Frequency of ICOS+ Tregs (% CD45) across MLN or spleen split per diet group. (e-g) Frequency of (e) Total RORγt expressing (CD4+ RORγt+), (f) RORγt+ FoxP3+ (Treg), (g) RORγt+ FoxP3- (Th17). P-values indicated, Kruskal-Wallis with post-hoc Dunn test for multiple comparisons. Coloured according to diet group: chow (orange), chow>western (yellow), western (dark blue) and western>chow (light blue). Dots indicate individual mice, bars indicate median. (h) Spearman’s correlation coefficients were calculated between relative abundance of significant genus level taxa identified using ANCOM-BC previously and the 6 Treg subsets (Supplementary Figure 7) across the (a) MLN. Multiple comparisons p-value correction was performed using the Benjamini-Hochberg method. Only significant correlations are shown (q <0.05). Colours correspond to correlation coefficients (numeric R values also indicated). + or - indicate the expression of RORγt and ICOS. Only Treg1-4 had significant correlations of R> 0.5 or R<0.5 and are presented.

**Supplementary Figure 7.**
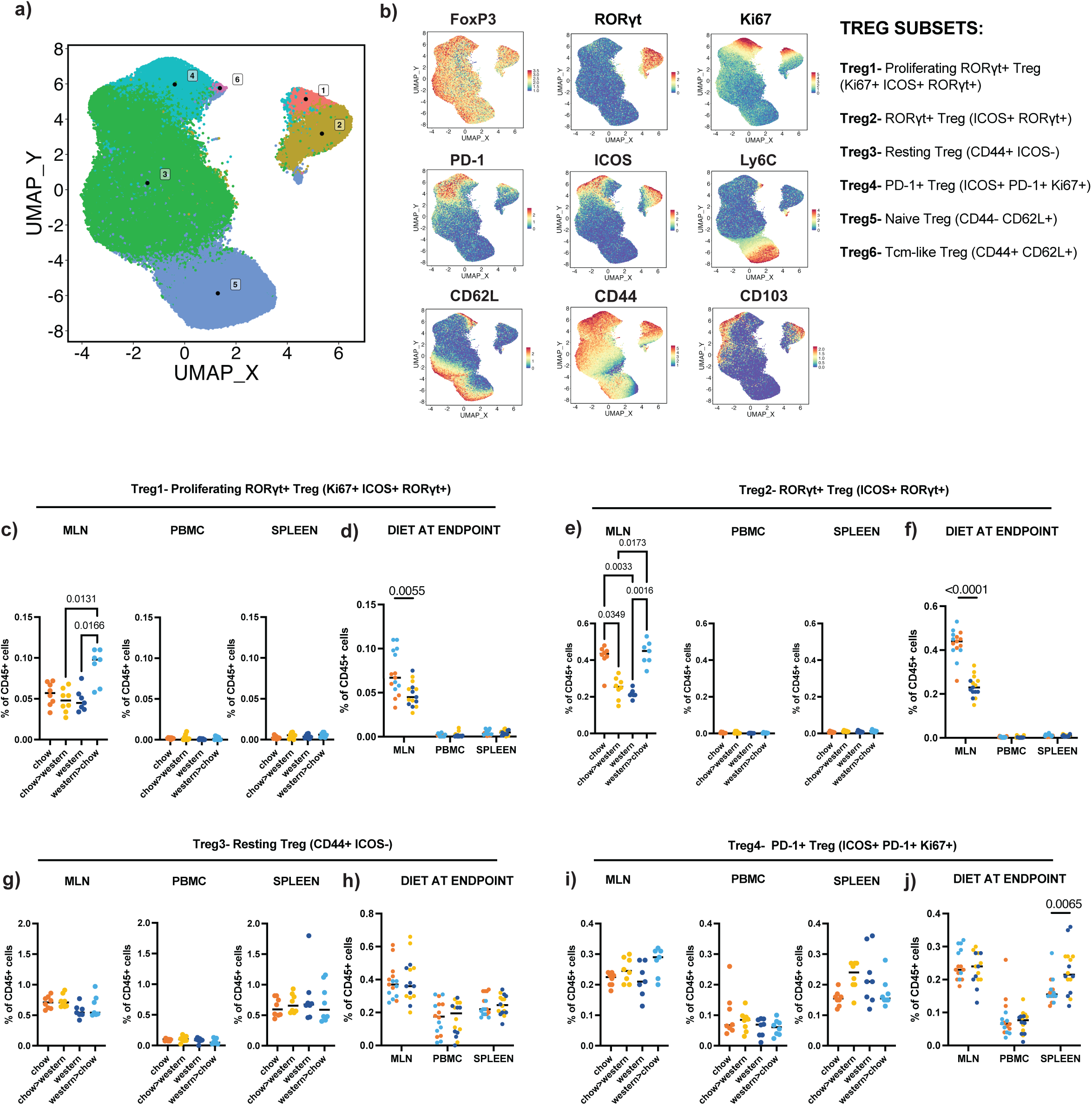
(a) Unsupervised FlowSOM clustering was performed on CD25+FoxP3+ cells per mouse across the MLN, PBMC and spleen to further interrogate the Treg compartment. FlowSOM clustering identified 6 metaclusters of Tregs (classified as Treg1-6). (a) UMAP plot visualising Treg subsets, coloured and annotated according to the metacluster (b) Expression of key markers defining Treg clusters overlayed onto UMAP, corresponding identity of Treg1-6 outlined in key. (c-j) To validate the clustering, Treg populations were manually gated. Frequency of (c-d) proliferating RORγt+ Treg (Ki67+ ICOS+ RORγt+) (Treg1), (e-f) RORγt+ Treg (ICOS+ RORγt+) (Treg2), (g-h) resting Tregs (CD44+ ICOS-) or (i-j) PD-1+ Tregs (ICOS+ PD-1+ Ki67+) as a percentage of CD45+ cells across MLN, PBMC or spleen, split per diet group or pooled according to diet at the endpoint. P-values indicated, Mann-Whitney U rank or Kruskal-Wallis with post-hoc Dunn test for multiple comparisons presented where appropriate. Coloured according to diet group: chow (orange), chow>western (yellow), western (dark blue) and western>chow (light blue). Dots indicate individual mice, bars indicate median.

